# Investigating the outcomes of virus coinfection within and across host species

**DOI:** 10.1101/2022.12.05.519092

**Authors:** Ryan M. Imrie, Sarah K. Walsh, Katherine E Roberts, Joanne Lello, Ben Longdon

## Abstract

Interactions between coinfecting pathogens have the potential to alter the course of infection and can act as a source of phenotypic variation in susceptibility between hosts. This phenotypic variation may influence the evolution of host-pathogen interactions within host species and interfere with patterns in the outcomes of infection across host species. Here, we examine experimental coinfections of two *Cripaviruses* – Cricket Paralysis Virus (CrPV), and Drosophila C Virus (DCV) –across a panel of 25 *Drosophila melanogaster* inbred lines and 47 *Drosophilidae* host species. We find that interactions between these viruses alter viral loads across *D. melanogaster* genotypes, with a ~3 fold increase in the viral load of DCV and a ~2.5 fold decrease in CrPV in coinfection compared to single infection, but we find little evidence of a host genetic basis for these effects. Across host species, we find no evidence of systematic changes in susceptibility during coinfection, with no interaction between DCV and CrPV detected in the majority of host species. These results suggest that phenotypic variation in coinfection interactions within host species can occur independently of natural host genetic variation in susceptibility, and that patterns of susceptibility across host species to single infections can be robust to the added complexity of coinfection.

## Introduction

Coinfections – simultaneous infections of a host with multiple pathogen lineages or species – are ubiquitous in nature, and represent the real-world context in which many infections occur [1–3]. Interactions between pathogens during coinfection can alter the virulence experienced by the host, and the loads and transmission rates of one or both pathogens [4–9]. At a population level, these interactions can lead to changes in infectious disease dynamics [10,11], such as the exclusion of novel viruses from host populations with other established pathogens [12,13], or fluctuations in the epidemic spread of one virus depending on the prevalence of other viruses [14,15]. These changes can ultimately alter the selective pressures imposed on hosts and pathogens, and coinfections have been proposed as a mechanism for the maintenance of genetic diversity in pathogen populations; as the fitness of pathogen genotypes may fluctuate not only in red queen dynamics with the host but also with coinfection prevalence and a pathogen’s competitive ability across coinfection scenarios [16]. Despite this, coinfections remain a largely understudied source of phenotypic variation during infection, and further investigation of their influence on the outcome of infection in different hosts and host species is needed.

Within coinfected hosts, pathogens can interact directly, such as through the production of toxins or modulation of the opposing pathogen’s gene expression [17,18], or indirectly through the production of common goods, competition for host resources, and interactions with host gene expression and immunity [19–24]. For example, in HIV-virus coinfections – a mechanistically well studied set of interactions due to their suspected involvement in AIDS progression [25] – several viruses have been shown to alter susceptibility to HIV infection by changing the expression of cell surface receptors CD4 and CCR5 [26,27]. In the case of human cytomegalovirus (HCMV), which upregulates CCR5 expression and increases HIV viral loads in coinfected tissues, HIV can reciprocally induce the expression of transmembrane proteins that promote HCMV infection [28,29]. Conversely, measles virus coinfections can inhibit HIV-1 replication due to measles-related activation of proinflammatory cytokines [30]. As such, the presence of coinfecting viruses may enhance or interfere with the ability of a virus to effectively establish an infection in a host, with these interactions often mediated by host components.

Despite the known role of host components in many coinfections, the extent to which host genetic variation in these components can influence the strength of interactions between pathogens – and so the ability of hosts to evolve directly to selective pressures imposed by coinfection – is unknown. Evidence suggesting a role of host genetics in the outcomes of coinfections is limited; however, several studies in plants have shown that pathogen community composition, coinfection prevalence, and disease severity during coinfection can vary non-randomly between host genotypes [31–33]. Coinfections can also be influenced by host dietary choices and the quantity of nutrients available in the host [34,35] – both of which are heritable traits [36,37] – which suggests that host genetic variation may influence coinfection outcomes. Broadly, we may expect host genetic variation to lead to changes in the strength of interaction between coinfecting pathogens when the interaction occurs through modulation of a host component (e.g., immune pathways or resource competition), or when host genetic variation influences the pathogen load of one or both pathogens.

Variation in the outcomes of coinfection across host species has also received relatively little attention, with most comparative cross-species studies focusing either on single infections in controlled experimental systems [38–45] or looking for broad patterns in infections across large datasets of natural systems where coinfection status is unknown [46–52]. These studies have shown that the evolutionary relationships between host species can explain a large proportion of the variation in infection traits. For example, virulence tends to increase [45–47], and onward transmission and pathogen load decrease [39,46], with greater evolutionary distance between donor and recipient hosts. Irrespective of distance to the donor host, closely related species also tend to share similar levels of susceptibility to novel pathogens [39–41]. Phylogenetic models such as these form part of the growing field of zoonotic risk prediction, the aim of which is to provide accurate, actionable predictions of host-virus interactions to inform public health measures [53]. The accuracy of current models may be improved by identifying and incorporating additional sources of variation in the outcome of cross-species transmission [54]. Coinfection status, detectable through metagenomic screening [55], may be a beneficial inclusion in such models, provided the strength and/or direction of coinfection interactions are known (or inferable).

Here, we investigate the influence of coinfection on virus susceptibility within and across host species, using panels of *Drosophila* hosts and experimental infections with two

### Cripaviruses

Cricket Paralysis Virus (CrPV) and Drosophila C Virus (DCV). Viral loads were measured during single and coinfection conditions across 25 inbred lines of *Drosophila melanogaster* and 47 *Drosophilidae* species. By analysing both viral loads and the change in viral loads from single to coinfection, we quantify the host genetic and phylogenetic components of susceptibility to each virus, and investigate whether these host components also influence the strength and direction of coinfection interactions in this system.

Both DCV and CrPV are well studied pathogens in *Drosophila melanogaster* and multiple similarities exist in their interactions with their hosts that could lead to interactions during coinfection. Both viruses are targeted by the antiviral RNAi pathway during infection of *D. melanogaster* [56,57], and activate the IMD immune signalling pathway, inducing non-specific antiviral gene expression [58–60]. Each encodes an inhibitor of antiviral RNAi, which act on different components of the pathway; the DCV inhibitor binds and sequesters viral RNA to prevent its cleavage by the antiviral RNAi endonuclease *Dicer-2*, and also disrupts formation of the RNA-induced silencing complex (RISC) [61,62]; the CrPV inhibitor binds the RISC protein *Argonaute-2*, causing suppression of RISC viral RNA cleavage [62]. Infections with DCV have also been shown to induce nutritional stress in infected hosts, due to intestinal obstruction and accumulation of food in the fly crop, although CrPV infection results in no such phenotype [63]. DCV and CrPV may therefore be capable of interacting indirectly during coinfection through multiple routes: by suppression of antiviral RNAi, transactivation of host antiviral gene expression, or competition for limited host resources.

Susceptibility to DCV infection has a strong host genetic component [64], with polymorphisms in two major-effect genes (*pastrel* and *Ubc-E2H*) explaining a large proportion of the variation in DCV susceptibility [64–67]. Both genes have also been implicated in CrPV susceptibility during knockdown experiments [67]. DCV and CrPV both vary widely in their ability to persist and replicate across different *Drosophilidae* host species, with the host phylogeny explaining a large proportion of the variation in viral load during single infection [40,41]. A role of host genetics during coinfection may therefore manifest either as a change in the genetic/phylogenetic components of susceptibility to each individual virus, or as a genetic/phylogenetic component directly influencing the strength of interaction between these viruses, both of which we investigate here.

## Materials & Methods

### Fly stocks

Stocks of DGRP flies were kindly provided by Jon Day and Francis Jiggins [64]. In total, 25 DGRP lines were used (for details see Supplementary Table 1), with 15 lines containing the resistant “G” allele of the A2469G *pastrel* SNP and 10 containing the susceptible “A” allele [65].*Pastrel* allele status was confirmed via conventional PCR using SNP genotyping primers from [65] (Supplementary Table 2). Laboratory stocks of 47 *Drosophilidae* host species were used to provide the across-species host panel (Supplementary Table 3), as in previous studies [40,41].

All flies were maintained in multi-generation stock bottles (Fisherbrand) at 22°C, 70% relative humidity in a 12-hour light-dark cycle. Each stock bottle contained 50ml of one of four varieties of *Drosophila* media (https://doi.org/10.6084/m9.figshare.21590724.v1) which were chosen to optimise rearing conditions for parental flies. All fly lines and species were confirmed to be negative for infection with CrPV and DCV prior to experiments by quantitative reverse-transcription PCR (qRT-PCR, described below). To limit the effects of variation in larval density on the condition of DGRP lines, experimental flies were reared in vials with finite numbers of larvae, achieved by transferring groups of five 7 day old mated females to fresh vials each day for 3 days, with daily pools of offspring from these vials collected for experiments. Due to large differences in fecundity, larval density controls were not practical for the across-species host panel.

### Inferring the Drosophilidae host phylogeny

The method used to infer the host phylogeny has been described in detail elsewhere [40]. Briefly, publicly available sequences of the *28S, Adh, Amyrel, COI, COII, RpL32*, and *SOD* genes were collected from Genbank (see https://doi.org/10.6084/m9.figshare.13079366.v1 for a full breakdown of genes and accessions by species). Gene sequences were aligned in Geneious version 9.1.8 (https://www.geneious.com) using a progressive pairwise global alignment algorithm with free end gaps and a 70% similarity IUB cost matrix. Gap open penalties, gap extension penalties, and refinement iterations were kept as default.

Phylogenetic reconstruction was performed using BEAST version 1.10.4 [68], as the subsequent phylogenetic mixed model (described below) requires a tree with the same root-tip distances for all taxa. Genes were partitioned into separate ribosomal (*28S*), mitochondrial (*COI, COII*), and nuclear (*Adh*, *Amyrel, RpL32, SOD*) groups. The mitochondrial and nuclear groups were further partitioned into groups for codon position 1+2 and codon position 3, with unlinked substitution rates and base frequencies across codon positions. Each group was fitted to separate relaxed uncorrelated lognormal molecular clock models using random starting trees and four-category gamma-distributed HKY substitution models. The BEAST analysis was run twice, with 1 billion Markov chain Monte Carlo (MCMC) generations sampled every 100,000 iterations, using a birth-death process tree-shape prior. Model trace files were evaluated for chain convergence, sampling, and autocorrelation using Tracer version 1.7.1 [69]. A maximum clade credibility tree was inferred from the posterior sample with a 10% burn-in. The reconstructed tree was visualised using ggtree version 2.0.4 [70].

### Virus isolates

Virus stocks were kindly provided by Julien Martinez (DCV) [71], and Valérie Dorey and Maria Carla Saleh (CrPV) [61]. The DCV isolate used here (DCV-C) was isolated from lab stocks established by wild capture in Charolles, France [72], and the CrPV isolate was collected from *Teleogryllus commodus* in Victoria, Australia [73]. Virus stocks were checked for contamination with CrPV (DCV) and DCV (CrPV) by qRT-PCR and diluted in Ringers solution [74] to equalise the relative concentrations of viral RNA. Before inoculation, virus aliquots were either mixed 1:1 with Ringers (single infection inoculum) or 1:1 with an aliquot of the other virus (coinfection inoculum). This was done to keep the individual doses of each virus consistent between infection conditions.

### Inoculation

Before inoculation, 0-1 day old male flies were transferred to vials containing cornmeal media. These flies were then transferred to fresh media every 2 days for a week (age 7-8 days), at which point they were inoculated. Vials contained between 7 and 20 flies (mean = 12.7), and were kept at 22°C, 70% relative humidity in a 12-hour light-dark cycle throughout the experiments. Male flies were used to avoid any effect of sex or mating status, which has been shown to influence the susceptibility of female flies to other pathogens [75–77]. Flies were inoculated under CO_2_ anaesthesia via septic pin prick with 12.5μm diameter stainless steel needles (Fine Science Tools, CA, USA). These needles were bent approximately 250μm from the end to provide a depth stop and dipped in virus inoculum before being pricked into the pleural suture of anaesthetised flies. Inoculation by this method bypasses the gut immune barrier but avoids differences in inoculation dose due to variation in feeding rate, and infections using this route largely follow the same course as oral infections but with less stochasticity [78].

### Measuring change in viral load

To provide a measure of viral load during single and coinfection, inoculated flies were snap frozen in liquid nitrogen at 2 days (± 2 hours) post-inoculation. Additional samples were collected for each species in the *Drosophilidae* host panel immediately after inoculation, which were used to account for differences in housekeeping gene expression across host species during C_T_ normalisation (see below). Total RNA was extracted from flies homogenized in Trizol (Invitrogen) using chloroform-isopropanol extraction, and reverse transcribed using GoScript reverse transcriptase (Promega) with random hexamer primers. qRT-PCR was carried out on 1:2 diluted cDNA on an Applied Biosystems StepOnePlus system using a Sensifast Hi-Rox SYBR kit (Bioline). Cycle conditions were as follows: initial denaturation at 95°C for 120 seconds, then 40 cycles of 95°C for 5 seconds and 60°C for 30 seconds. The primer pairs used for virus qRT-PCR assays were: (DCV) forward, 5’-GACACTGCCTTTGATTAG-3’; reverse, 5’-CCCTCTGGGAACTAAATG-3’; (CrPV) forward, 5’-TTGGCGTGGTAGTATGCGTAT-3’; reverse, 5’-TGTTCCGTCCTGCGTCTC. RPL32 housekeeping gene primers were used for normalisation and varied by species (Supplementary Table 4-5). For each biological sample, two technical replicate qRT-PCR reactions were performed for each amplicon (viral and RPL32).

Between-plate variation in C_T_ values was estimated and corrected using linear models with plate ID and biological replicate ID as fixed-effects [79,80]. For DGRP lines, mean viral C_T_ values from technical replicate pairs were normalised to RPL32 and converted to relative viral load using the ΔΔCτmethod, where ΔC_T_ = C_T:Virus_ – C_T:RPL32_ and ΔΔC_T_ = 40 – ΔC_T_. To account for potential differences in RPL32 expression between species, change in viral load in the *Drosophilidae* species experiment was calculated as fold-change in viral load from inoculation to 2 days post-infection using the ΔΔC_T_ method, where ΔC_T_ = C_T:Virus_ – C_T:RPL32_ and ΔΔC_T_ = ΔC_T:day0_ – ΔC_T:day2_. Amplification of the correct products was verified by melt curve analysis. Repeated failure to amplify product, the presence of melt curve contaminants, or departures from the melt curve peaks of positive samples (±1.5°C for viral amplicons; ±3°C for RPL32) were used as exclusion criteria for biological replicates. For a full breakdown of the replicates per experiment for each combination of fly line/species and infection condition see Supplementary Table 6.

### Analysis of coinfection within and across species

Genetic variation in the outcome of single and coinfection across DGRP lines was analysed using methods previously described by Magwire et al. [64]. Briefly, multivariate generalised linear mixed models (GLMMs) were fitted using the R package MCMCglmm [81], with either the viral loads of each virus under each infection condition, or the change in viral load during coinfection (coinfection viral load - single infection viral load) as the response variable. The structures of these models were as follows:

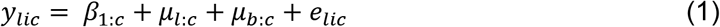

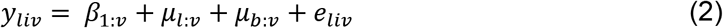

In model (1), *y_lic_* is the viral load for the combination of virus and infection condition *c* (CrPV single infection, CrPV coinfection, DCV single infection, DCV coinfection) in the *i^th^* biological replicate of DGRP line *l*. The fixed effect *β_1_* represents the intercepts for each combination, the random effect *μ_l_* represents the deviation of each DGRP line from the overall mean viral load for each combination (equivalent to the between-line variance), and *e_lic_* represents the residual error. A small but significant effect of experiment block was found in initial models, driven by ~10 fold differences in DCV viral loads of the third experimental block. To account for this, random effects of block by infection condition (*μ_b:c_, μ_b:v_*) were added to both models. The structure of model (2) remains the same, but with the change in viral load during coinfection for each virus as the response variable, and *y_liv_* representing the change in viral load for the *i^th^* biological replicate of virus *v* and DGRP line *l. Pastrel* allele status (susceptible “A”, resistant “G”) was included in additional models as a fixed effect (*β_2:lp_*).

Phylogenetic GLMMs were used to investigate the effects of host evolutionary relatedness on viral load during single and coinfection, and to calculate interspecific correlations between different infection conditions across host species. Multivariate models were fitted with the viral loads of each virus under each infection condition as the response variable. The structure of these models were as follows:

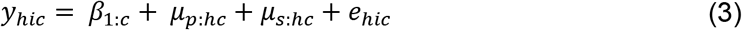

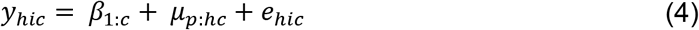

In these models, *y_hic_* is the change in viral load for the combination of virus and infection condition *c* (CrPV single infection, CrPV coinfection, DCV single infection, or DCV coinfection) in the *i^th^* biological replicate of host species *h*. The fixed effect *β_1_* represents the intercepts for each combination, the random effect *μ_p_* represents the effects of the host phylogeny assuming a Brownian motion model of evolution, and *e* represents the model residuals. Model (3) also includes a species-specific random effect that is independent of the host phylogeny (*μ_s:hc_*). This explicitly estimates the non-phylogenetic component of between-species variance and allows the proportion of variance explained by the host phylogeny to be calculated. *μ_s:hc_* was removed from model (4) as model (3) struggled to separate the phylogenetic and species-specific traits for some infection conditions. Wing size, measured as the length of the IV longitudinal vein from the tip of the proximal segment to the join of the distal segment with vein V [82], provided a proxy for body size [83] and was included in a further model as a fixed effect (*wingsizeβ_2:hc_*). This was done to ensure that any phylogenetic signal in body size did not explain the differences seen in viral load between species [84].

To investigate the effect of host evolutionary relatedness on the change in viral load from single to coinfection, additional models were run with the change in viral load during coinfection (coinfection viral load - single infection viral load) on viral load as the response variable:

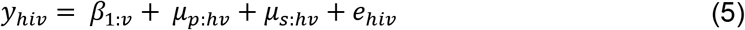

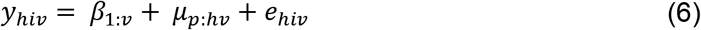

In these models, *y_hiv_* is the change in viral load for the *i^th^* biological replicate of virus *v* and host species *h*. The explanatory structure otherwise remains the same as models (3-4).

Within models (1-6), the random effects and residuals were assumed to follow a multivariate normal distribution and a centred mean of 0. Models (1-2) were fitted with a covariance structure *V_t_*⊗*l* for the between line variances, and *V_e_* ⊗ *I* for the residuals, with ⊗ representing the Kronecker product, and *I* representing an identity matrix. *V* represents 4 x 4 covariance matrices for model (1) and 2 x 2 covariance matrices for model (2) which describe the between-line variances and covariances in viral load for each infection condition and virus. Models (3-6) were fitted with a covariance structure of *V_p_* ⊗ *A* for the phylogenetic effects, *V_s_* ⊗ *I* for species-specific effects, and *V_e_* ⊗ *I* for residuals. *A* represents the host phylogenetic relatedness matrix, *I* an identity matrix, and *V* represents 4 × 4 covariance matrices for models (3-4), or 2 x 2 covariance matrices for models (5-6), describing the between-species variances and covariances of changes in viral load for each combination of virus and infection condition. As each biological replicate was only tested with one combination of virus and infection condition, the covariances of *V_e_* cannot be estimated and were set to 0 for all models.

Models were run for 13 million MCMC iterations, sampled every 5000 iterations with a burn-in of 3 million iterations. Parameter expanded priors were placed on the covariance matrices, resulting in multivariate F distributions with marginal variance distributions scaled by 1000. Inverse-gamma priors were placed on the residual variances, with a shape and scale equal to 0.002. To ensure the model outputs were robust to changes in prior distribution, models were also fitted with flat and inverse-Wishart priors, which gave qualitatively similar results. All parameter estimates reported from models (1-6) are means of the posterior density, and 95% credible intervals (CIs) are the 95% highest posterior density intervals which are reported in brackets following the estimates in the results.

The covariance matrices of models (1) and (2) were used to calculate the heritabilities (*h^2^*), and covariates of additive genetic and environmental variation (*CV_A_* and *CV_E_* respectively) of viral load and the effects of coinfection within host species. Heritability was calculated as 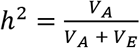, where *V_A_* represents the additive genetic variance and *V_E_* the environmenta variance of each trait [85]. As DGRP lines are homozygous, *V_A_* can be calculated as half the between-line variance, assuming purely additive genetic variation [64]. *V_E_* was set as the residual variance of each model, which contains both non-additive genetic and environmental effects on viral load and any measurement errors. Genetic correlations between infection conditions were calculated from the model (1) and (2) *v_l_* matrices as 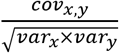 and slopes of each relationship as 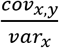

The proportion of the between species variance that can be explained by the phylogeny was calculated from models (3) and (5) using the equation 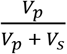, where *V_p_* and *v_s_* represent the phylogenetic and species-specific components of between-species variance respectively [84], and are equivalent to phylogenetic heritability or Pagel’s lambda [86,87]. The repeatability of viral load measurements was calculated from models (4) and (6) as 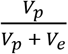 where *Ve* is the residual variance of the model [88]. Interspecific correlations in viral load between single and coinfection were calculated from model (4) *V_p_* matrix as, 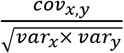.

### Data Availability

Data and R scripts for all included statistical models can be found at https://doi.org/10.6084/m9.figshare.21657503.v1.

## Results

### Coinfection causes changes in DCV and CrPV viral load across D. melanogaster genotypes

To investigate variation in the outcome of coinfection within host species, we injected a total of 8,618 flies from 25 lines of the Drosophila Genetic Reference Panel with one of three virus inoculums: DCV, CrPV, and DCV + CrPV, and measured the outcome of infection as the viral load of each virus at 2 days post-inoculation using qRT-PCR (Fig. 1). Point estimates of the mean viral load across lines suggest that DCV viral load increases ~3-fold during coinfection with CrPV, and CrPV viral load decreases ~2.5-fold during coinfection with DCV, although credible intervals of these estimates overlapped (Table 1). When models were fitted on the change in viral load (coinfection - single infection), in effect treating viral loads within experiment blocks as paired data, similar and significant effects of coinfection across lines were detected (Table 1). Several lines showed notably large changes during coinfection: two DGRP lines showed ~10 fold decreases in CrPV viral load, and three lines showed ~40-150 fold increases in DCV viral load. Removing these lines from model (2) reduced the mean changes in viral load during coinfection to a ~2 fold increase for DCV and a ~2 fold decrease for CrPV, but the effects of coinfection on both viruses remained significant.

**Figure 1:**
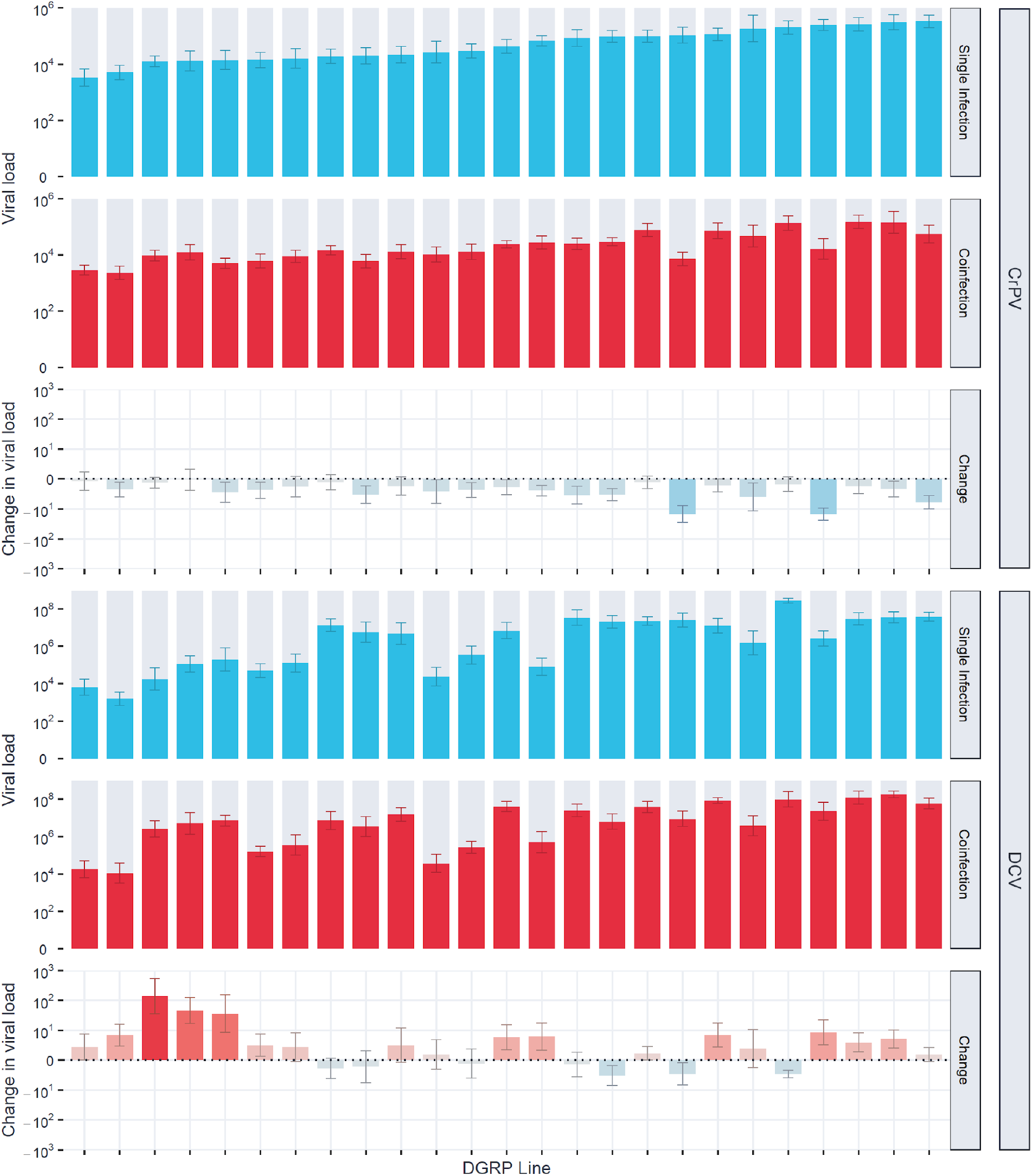
Viral loads of CrPV and DCV across DGRP lines during single and coinfection. Bar heights show the mean viral load or changes in viral load (coinfection - single infection) at 2 dpi on a log_10_ scale, with error bars showing the standard error of the mean. Blue bars represent single infection viral loads, or changes in viral load where single infection viral loads were greater than coinfection viral loads. Red bars represent coinfection viral loads, or changes in viral load where coinfection viral loads were greater than single infection viral loads. DGRP lines are arranged on the x-axis in order of susceptibility to CrPV during single infection.

**Table 1:** Estimates of the phenotypic mean, environmental variance (*V_E_*), additive genetic variance (*V_A_*), and heritability (*h^2^*) of viral load and the change in viral load during coinfection across DGRP lines for CrPV and DCV during single infection and coinfection. Values for “single infection” and “coinfection” conditions were taken from model (1), which was fitted on log_10_-transformed fold-changes in viral load, while values for “change” were taken from model (2), which was fitted on log10-transformed Δ fold-changes in viral load (coinfection - single infection).

| **Virus** | **Condition** | **Mean** | **$V_E$** | **$V_A$** | **$h^2$** |
| --- | --- | --- | --- | --- | --- |
| **CrPV** | Single Infection | 4.68 (4.02, 5.34) | 0.94 (0.79, 1.09) | 0.14 (0.05, 0.25) | 0.13 (0.05, 0.22) |
|  | Coinfection | 4.28 (3.60, 4.96) | 0.74 (0.61, 0.86) | 0.12 (0.04, 0.20) | 0.13 (0.06, 0.22) |
|  | Change | -0.27 (-0.37, -0.18) | 0.23 (0.15, 0.27) | 0.00 (0.00, 0.02) | 0.02 (0.00, 0.09) |
| **DCV** | Single Infection | 6.17 (5.29, 6.98) | 2.08 (1.75, 2.46) | 1.00 (0.47, 1.65) | 0.32 (0.20, 0.46) |
|  | Coinfection | 6.62 (5.82, 7.40) | 1.80 (1.49, 2.09) | 0.71 (0.33, 1.18) | 0.28 (0.16, 0.40) |
|  | Change | 0.33 (0.18, 0.47) | 0.21 (0.15, 0.27) | 0.03 (0.00, 0.06) | 0.11 (0.00, 0.23) |

### No evidence of a host genetic component to the outcome of coinfection

To estimate the influence of host genetic variation on the viral loads measured during single and coinfection, GLMMs were fitted to allow the phenotypic variation in viral loads to be partitioned into genetic and environmental components. Point estimates of the heritability of DCV viral load (0.25-0.30) were higher than for CrPV (0.13), and this difference was driven by changes in the genetic component of variation (Supplementary Table 7): CrPV CV_A_ = 0.08, (0.05, 0.11), DCV CV_A_ = 0.16, (0.12, 0.20). This is consistent with previous studies which also found the genetic component of variation in susceptibility of *D. melanogaster* to single infections with DCV (a natural pathogen) is higher than for CrPV (a novel pathogen) [64]. However, we found little evidence that heritability of DCV or CrPV viral loads change in relation to coinfection, with the credible intervals of *h^2^* estimates for single and coinfection viral loads overlapping for both viruses (Table 1). Additionally, no host genetic component was found for the change in viral load during coinfection (Table 1, Supplementary Table 8). Together, this suggests that variation in the strength of coinfection interactions between these viruses was independent of host genetic variation, and that coinfection status does not appear to alter the host genetic component of susceptibility to either virus.

Correspondingly, strong positive correlations between single and coinfection viral loads were found for both DCV: r = 0.94 (0.84, 1.00), and CrPV: r = 0.90 (0.73, 1.00), with little evidence of genotype-by-coinfection interactions. Strong positive correlations were also seen between the two viruses, such that DGRP lines more susceptible to DCV were often also more susceptible to CrPV (Fig 2). Together, this suggests that susceptibility to DCV and CrPV share similar genetic architectures within *D. melanogaster*, with host genetic variation affecting DCV viral load similarly affecting CrPV viral load, again irrespective of coinfection status.

**Figure 2:**
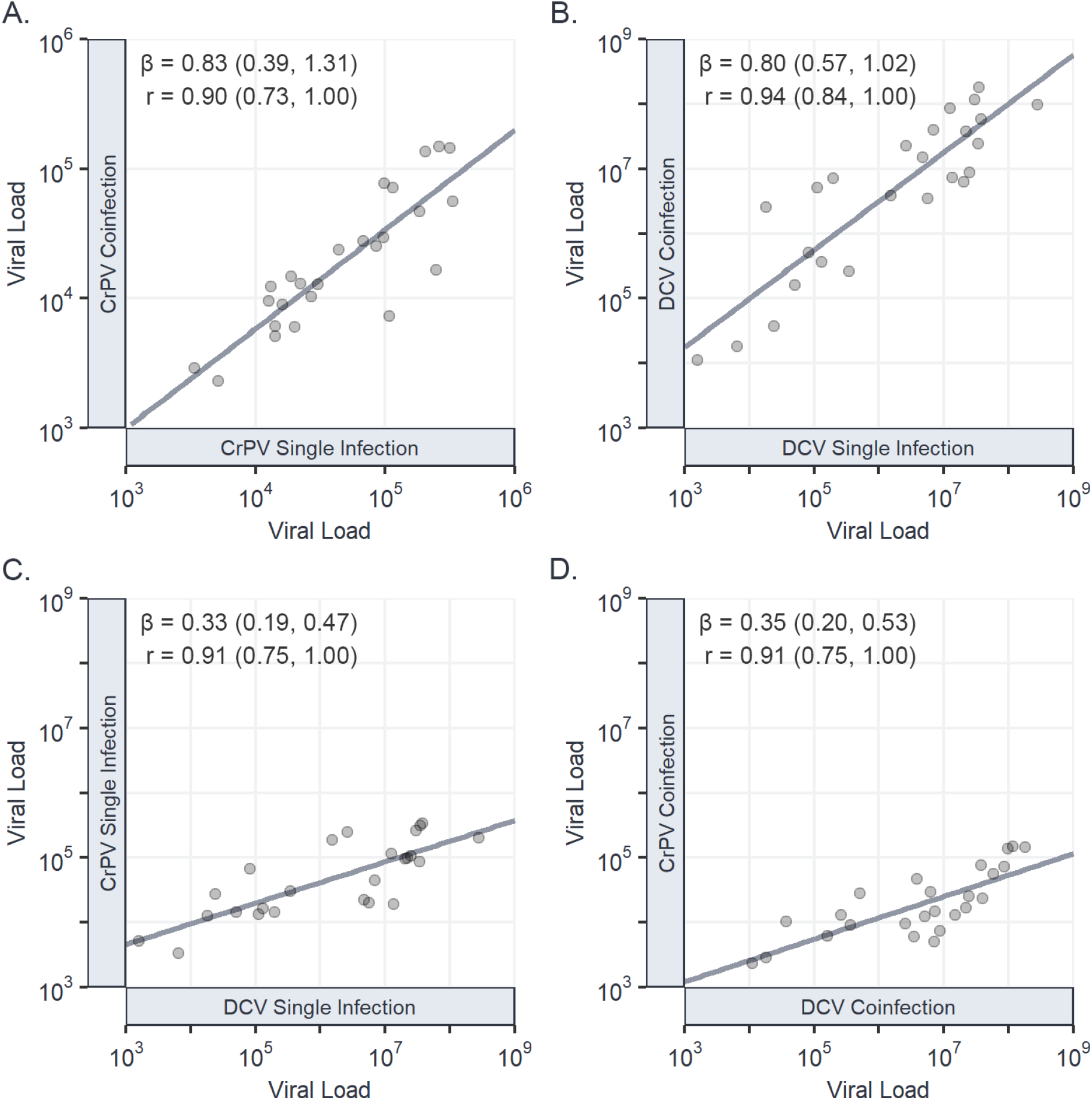
Genetic correlations in viral load between single and coinfections of CrPV and DCV. Correlations in viral load between CrPV during single and coinfection (A); DCV during single and coinfection (B); CrPV and DCV during single infection (C); and CrPV and DCV during coinfection (D). Individual points represent the mean viral load at 2 dpi for each DGRP line on a log10 scale, with trend lines added from a univariate least-squares linear model for illustrative purposes. Genetic correlations (r), regression slopes (β), and 95% CIs have been taken from the output of model (1).

### Viral load remains a repeatable trait across host species during coinfection

To investigate how coinfection may alter susceptibility across host species, we performed similar experimental single and coinfections across 47 *Drosophilidae* host species. A total of 13,596 flies were inoculated, and the change in viral load after two days of infection was measured by qRT-PCR (Fig 3). Neither virus showed evidence of changes in their overall mean viral loads or variance across host species between single and coinfection (Table 2). Power analysis based on the effects of coinfection found in *D. melanogaster* (Fig. 1, Supplementary Methods) showed that the level of replication in this experiment was adequate to detect systematic ~2-fold changes in viral load across host species. As such, this result suggests there is no evidence for large additive effects of coinfection that are consistent across host species. Instead, most host species showed no discernible differences in viral loads during coinfection, with notable exceptions including *D. obscura* (both viruses decreased in viral load by ~600 fold), *D. suzukii* (DCV unchanged but CrPV decreased by ~25 fold), *Zaprionus tuberculatus* (CrPV unchanged but DCV decreased by ~50 fold), and *D. virilis* (CrPV unchanged but DCV increased by ~40 fold).

**Figure 3:**
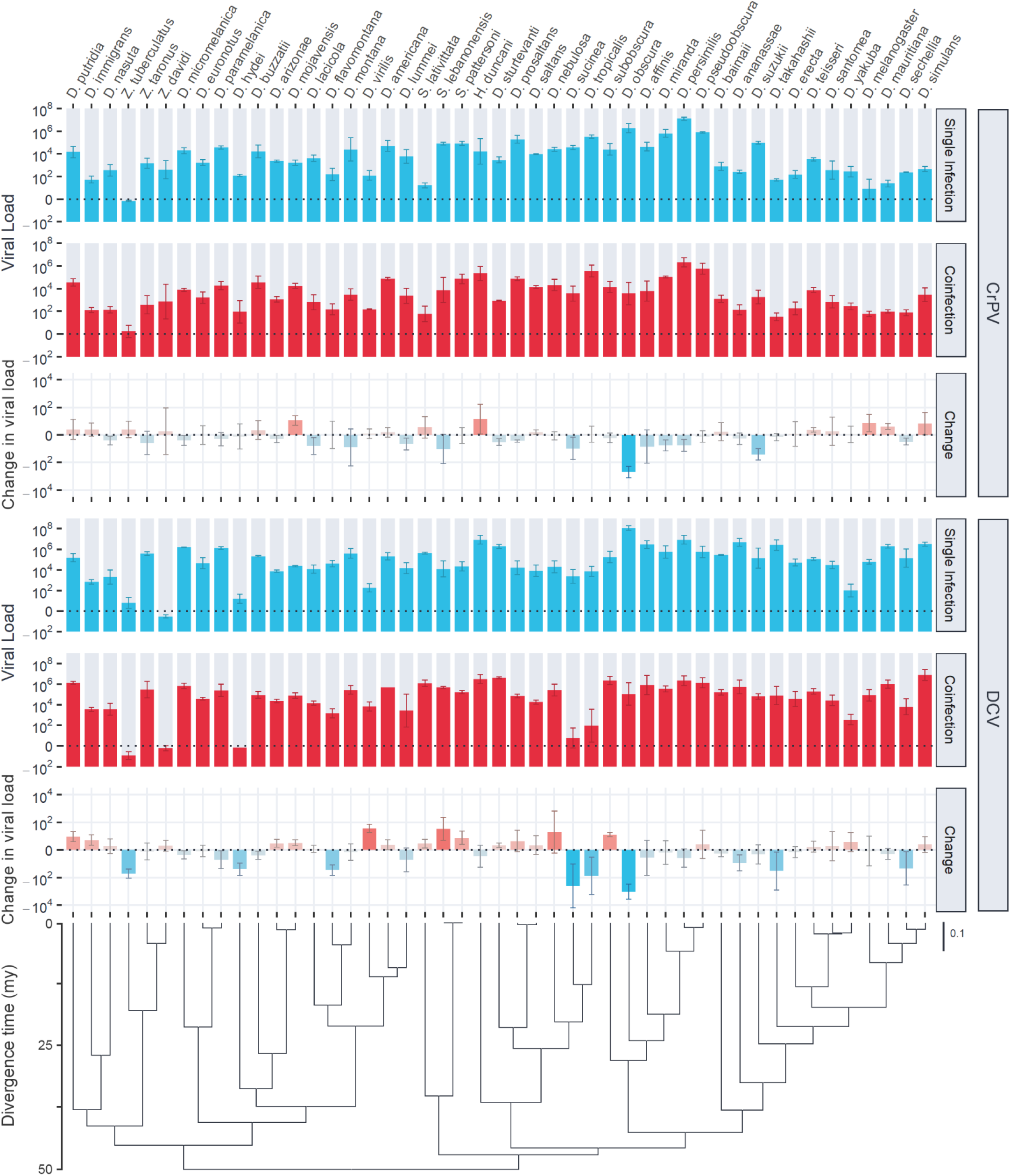
Viral loads of CrPV and DCV across host species during single and coinfection. Bar heights show the mean viral load or changes in viral load (coinfection – single infection) by 2 dpi on a log10 scale, with error bars showing the standard error of the mean. Blue bars represent single infection viral loads, or changes in viral load where single infection viral loads were greater than coinfection. Red bars represent coinfection viral loads, or changes in viral load where coinfection viral loads were greater than single infection. The phylogeny of *Drosophilidae* hosts is presented at the bottom, with the scale bar showing nucleotide substitutions per site, and the axis showing the approximate age since divergence in millions of years (mya) based on estimates from [89].

**Table 2:** Estimates of overall mean, across-species variance, repeatability, and the proportion of variance explained by the host phylogeny for viral load and the change in viral load during coinfection. Values for mean viral load, across species variance, and repeatability for the “single infection” and “coinfection” conditions were taken from model (4), which was fitted on log_10_-transformed fold-changes in viral load, while these values for “change” during coinfection were taken from model (6), which was fitted on log_10_-transformed Δ fold-changes in viral load (coinfection - single infection). The proportion of variance explained by phylogeny was taken from model (3) for the “single infection” and “coinfection” conditions, and model (5) for “change” during coinfection.

| Virus | Condition | Mean | Across-species variance | Repeatability | Variance explained by phylogeny |
| --- | --- | --- | --- | --- | --- |
| CrPV | Single Infection | 3.44 (1.88, 4.99) | 3.63 (1.86, 5.75) | 0.86 (0.78, 0.93) | 0.88 (0.69, 1) |
|  | Coinfection | 3.30 (2.21, 4.38) | 2.93 (1.36, 4.75) | 0.76 (0.64, 0.87) | 0.82 (0.59, 1) |
|  | Change | -0.13 (-0.49, 0.19) | 0.17 (0, 0.46) | 0.11 (0, 0.27) | 0.57 (0, 1) |
| DCV | Single Infection | 4.75 (2.09, 7.49) | 9.85 (5.20, 15.46) | 0.94 (0.90, 0.97) | 0.13 (0, 0.43) |
|  | Coinfection | 4.57 (1.82, 7.26) | 10.40 (4.87, 16.44) | 0.89 (0.82, 0.94) | 0.10 (0, 0.27) |
|  | Change | -0.12 (-1.08, 0.86) | 0.92 (0, 1.84) | 0.36 (0.08, 0.62) | 0.49 (0, 0.99) |

Phylogenetic GLMMs were fitted to the data to determine the proportion of variation in viral load explained by the host phylogeny (Table 2). The host phylogeny explained a large proportion of the variation in viral load for CrPV during single infection: 0.88 (0.69, 1), and coinfection: 0.82 (0.59, 1), with no credible difference between these two estimates. Estimates of the variation in DCV viral load explained by phylogeny were low: 0.1-0.13 with wide credible intervals due to model (3) struggling to separate phylogenetic and non-phylogenetic effects for DCV. The repeatability of viral load across host species was high for both viruses during single infection, CrPV: 0.86 (0.78, 0.93), DCV: 0.94 (0.90, 0.97) and coinfection, CrPV: 0.76 (0.64, 0.87), DCV: 0.89 (0.82, 0.94), with the between-species phylogenetic component explaining a high proportion of the variation in viral load with little within-species variation or measurement error. Although point estimates of these parameters were all consistent with a slight decrease in phylogenetic signal during coinfection, the effect size was small, and we did not detect credible differences in phylogenetic signal between single and coinfection.

### Viral load is strongly correlated between single and coinfection across host species

Interspecific correlations in viral load between single and coinfection were calculated for each virus from the variance-covariance matrix of model (4). We found strong positive correlations in viral loads between single and coinfection for DCV: r = 0.95 (0.89, 0.99) (Fig 4A) and CrPV: r = 0.94 (0.86, 0.99) (Fig 4B), with the regression slopes of each indicating a near 1:1 relationship: DCV: β = 0.98 (0.77, 1.22), CrPV: β = 0.85 (0.66, 1.05), and limited evidence of host species by coinfection interactions. The strength of the interspecific correlation in viral load between DCV and CrPV (Fig. 4C, D) also did not differ between single: r = 0.59 (0.31, 0.82), and coinfection: r = 0.67 (0.43, 0.88) and was consistent with previous estimates: r = 0.59 (0.26, 0.87) [41].

**Figure 4:**
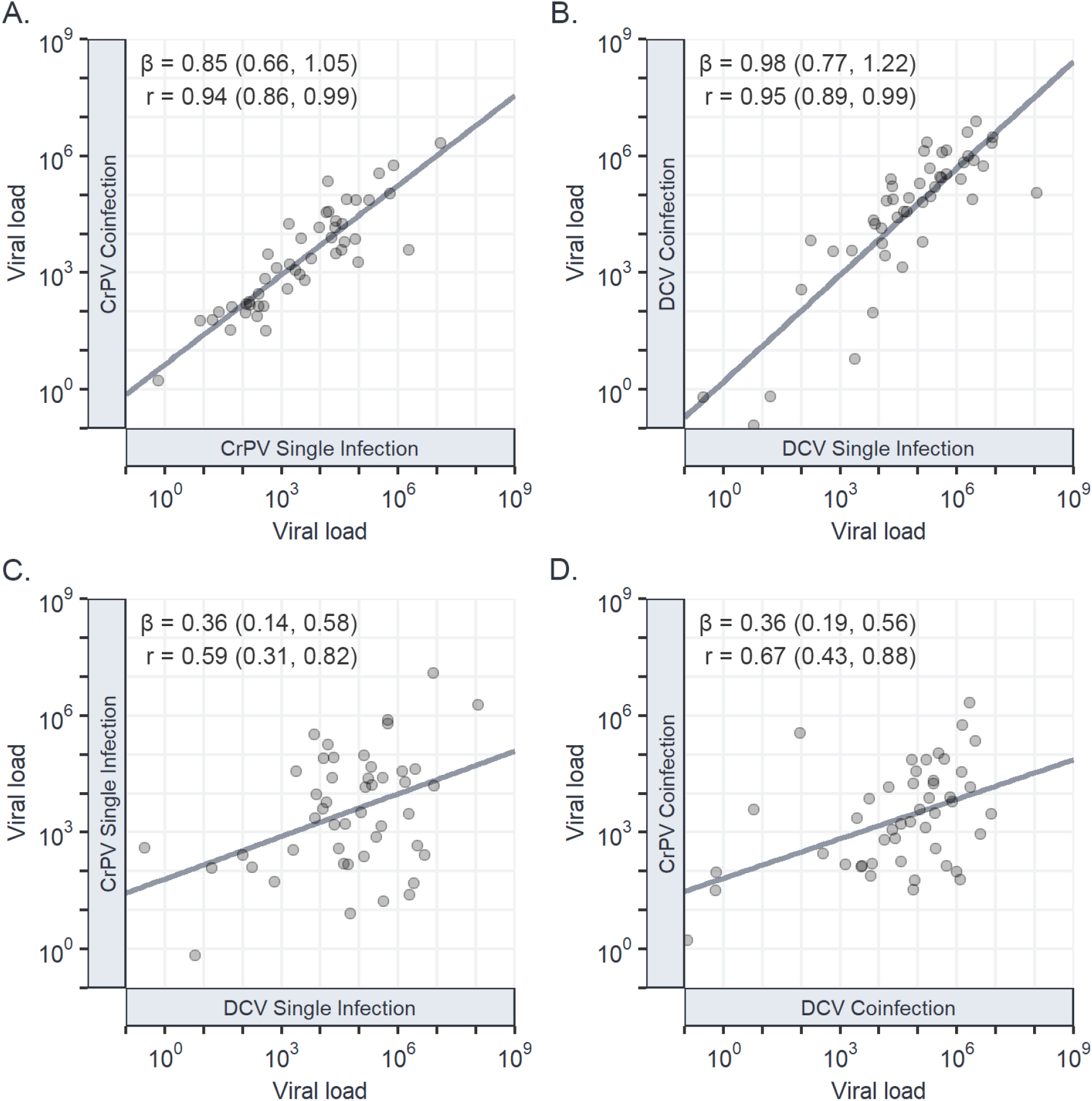
Interspecific correlations in viral load between single and coinfections of CrPV and DCV. Correlations in viral load between CrPV during single and coinfection (A); DCV during single and coinfection (B); CrPV and DCV during single infection (C); and CrPV and DCV during coinfection (D). Individual points represent the mean viral load at 2 dpi for each *Drosophilidae* host species on a log_10_ scale, with trend lines added from a univariate least-squares linear model for illustrative purposes. Interspecific correlations (r), regression slopes (β), and 95% CIs have been taken from the output of model (4).

### Little evidence of phylogenetic signal in the strength of coinfection interaction

As the viral loads of DCV and CrPV show a strong phylogenetic signal across host species, we also tested if there was phylogenetic signal across hosts in the change in viral load from single to coinfection (Table 2). Fitting phylogenetic mixed models to these data revealed little support for any phylogenetic signal in the change in viral load during coinfection, with low estimates of repeatability for DCV: 0.36 (0.08, 0.62) and no credible difference from zero for repeatability of CrPV or the variance explained by phylogeny for either virus.

## Discussion

Here, we measured variation in the outcome of coinfections within and across host species, using a *Drosophila* experimental system and two *Cripaviruses:* DCV and CrPV. We found effects of coinfection on viral load across genotypes of *D. melanogaster*, with DCV increasing ~3 fold and CrPV decreasing ~2 fold during coinfection. Consistent with previous studies, we found that host genetic variation explained a large proportion of variation in susceptibility to single infections [64], but little evidence was found for a change in this genetic component of susceptibility in the presence of a coinfecting virus, or for a host genetic component to the strength of interaction between these viruses. Across host species, we found no evidence of consistent coinfection interactions between these viruses and no change in the phylogenetic patterns of susceptibility to each virus during coinfection, although coinfection interactions were apparent in a subset of host species. Strong positive correlations between single and coinfection viral loads, and between DCV and CrPV both within and across host species suggest that similar genetic architectures are underlying susceptibility to these viruses, and that susceptibility is largely independent of coinfection status.

Exploitative coinfection interactions – where one pathogen benefits from coinfection to the detriment of the other – have been described in intestinal parasites of wood mice and wild rabbits [90,91], and in mixed-genotype *Pseudomonas* infections in plants [92]. The mechanisms underlying exploitative coinfection interactions are unknown but may be due to differences in the relative importance of specific interactions between pathogens and the host in overall susceptibility. Within *D. melanogaster*, our data suggest that the strength of interaction between DCV and CrPV is not virus density-dependent, as more susceptible host genotypes did not experience increased changes in viral load with coinfection compared to more resistant genotypes. This suggests that the coinfection interaction within *D. melanogaster* is unlikely to be caused by resource competition between DCV and CrPV, as susceptible hosts experienced >100-fold higher viral loads for both DCV and CrPV compared to more resistant hosts with no evidence of limited virus replication. DCV may instead be benefiting from increased suppression of antiviral RNAi due to expression of the CrPV immune inhibitor [62], while CrPV is hindered by the activation of other mechanisms of host immunity by DCV. However, complex direct virus-virus interactions have been described in multiple coinfections, and it is possible that DCV and CrPV are directly influencing each other’s expression or virion surface composition [93,94].

Across host species, the changes in viral load during coinfection were highly variable and show no consistent interaction between DCV and CrPV. Coupled with the fact we did not detect effects of genetic variation within host species or evolutionary relatedness across host species in the change in viral load during coinfection, our results suggest that natural levels of variation in host genetics have little impact on the strength of interaction between these viruses during coinfection. This contrasts with coinfection studies in other systems, which describe variation between host genotypes in pathogen community composition, coinfection prevalence, and disease severity during coinfection [31–33]. Mathematical models investigating stochasticity during coinfection have suggested that otherwise identical coinfections can have directionally different outcomes [95], and so it may be that any influences of host evolutionary relatedness and genotype are being masked by high stochasticity in the outcome of coinfection in this system. Alternatively, variation in the strength of coinfection interaction between host genotypes may be influenced by a small number of major-effect loci that are not dispersed phylogenetically, which these experiments were not designed to detect.

As inferential and epidemiological models of cross-species infections grow in complexity, they will continue to incorporate more non-genomic data which is known to influence the outcome of infection (e.g., [96]). Our findings suggest that coinfection will not be a necessary inclusion in models of every host-pathogen system, as the ability of the host phylogeny to explain variation in viral load across host species was largely unaffected during coinfection in this case. Despite this, coinfection is known to cause changes in infection traits in many systems [4–9,97–109], with consequences for pathogen spread and establishment in natural populations [10–15]. Few studies exist that describe pathogens that do not interact during coinfection [110] (although this may represent publication bias), and so the frequency of consequential coinfection interactions in nature is as yet unknown. It remains unclear if interactions between pathogens can be consistently predicted *a priori* from single infection data [111,112], or from pathogen and host genomic data [113]. In cases of direct interaction between pathogens, such as the binding and activation of endogenous HIV by herpes simplex virus proteins [93], differing outcomes in coinfection may be predictable through conventional tools for inferring protein-protein and protein-nucleotide binding [114,115]. However, where pathogens interact indirectly, such as through immune modulation or resource availability, it may be necessary to understand the extent of variation in these host factors that is required to influence the outcome of infection before inferring interactions between coinfecting pathogens.

Here, we have tested for variation in the outcome of coinfection within and across host species, and our findings suggest that host genetics may not influence coinfection interactions in all host-pathogen systems. This approach can now be expanded to a more diverse range of coinfecting pathogens, to look for effects of host genetic variation during other pathogen-pathogen interactions, to better understand the potential determinants of the outcome of coinfection interactions, and how these interactions may affect the evolution of host susceptibility.

## Acknowledgements

We would like to thank Camille Bonneaud and Ann Tate for useful discussion. Ben Longdon and Ryan M. Imrie are supported by a Sir Henry Dale Fellowship jointly funded by the Wellcome Trust and the Royal Society (grant no. 109356/Z/15/Z), and a studentship funded by the Natural Environment Research Council (NERC) GW4+ Doctoral Training Partnership. Sarah K. Walsh is funded by a studentship from the Biotechnology and Biological Sciences Research Council (BBSRC) South West Biosciences Doctoral Training Partnership (BB/M009122/1). For the purpose of Open Access, the author has applied a CC BY public copyright licence to any Author Accepted Manuscript version arising from this submission.

## Notes

### Competing Interest Statement

The authors have declared no competing interest.

